# Engineering a culturable *Serratia symbiotica* strain for aphid paratransgenesis

**DOI:** 10.1101/2020.09.15.299214

**Authors:** Katherine M. Elston, Julie Perreau, Gerald P. Maeda, Nancy A. Moran, Jeffrey E. Barrick

## Abstract

Aphids are global agricultural pests and important models for bacterial symbiosis. To date, none of the native symbionts of aphids have been genetically manipulated, which limits our understanding of how they interact with their hosts. *Serratia symbiotica* CWBI-2.3^T^ is a culturable, gut-associated bacterium isolated from the black bean aphid. Closely related *Serratia symbiotica* strains are facultative aphid endosymbionts that are vertically transmitted from mother to offspring during embryogenesis. We demonstrate that CWBI-2.3^T^ can be genetically engineered using a variety of techniques, plasmids, and gene expression parts. Then, we use fluorescent protein expression to track the dynamics with which CWBI-2.3^T^ colonizes the guts of multiple aphid species, and we measure how this bacterium affects aphid fitness. Finally, we show that we can induce heterologous gene expression from engineered CWBI-2.3^T^ in living aphids. These results inform the development of CWBI-2.3^T^ for aphid paratransgenesis, which could be used to study aphid biology and enable future agricultural technologies.

**IMPORTANCE:** Insects have remarkably diverse and integral roles in global ecosystems. Many harbor symbiotic bacteria, but very few of these bacteria have been genetically engineered. Aphids are major agricultural pests and an important model system for the study of symbiosis. This work describes methods for engineering a culturable aphid symbiont, *Serratia symbiotica* CWBI-2.3^T^. These approaches and genetic tools could be used in the future to implement new paradigms for the biological study and control of aphids.

## INTRODUCTION

Many insects have a characteristic bacterial microbiome. These associations can take many different forms, ranging from the conserved gut communities of bees (1) to symbionts of sap-sucking insects that have evolved to resemble organelles (2). These and other bacterial-host relationships have inspired attempts to study and control insects by genetically engineering their resident microbiomes. This approach, known as “paratransgenesis”, has been developed primarily for insects that are vectors of human disease, including kissing bugs, tsetse flies, and mosquitoes (3–5). Recently, feasibility of paratransgenesis has also been demonstrated in agricultural pests, where it could provide an alternative to chemical pesticides and the development of genetically engineered crops (6–9).

Aphids are major worldwide agricultural pests and vectors for many plant viruses (10–12). They are also model organisms for understanding insect-endosymbiont coevolution because they have evolved close relationships with multiple species of bacterial symbionts. Typically, aphid symbionts are housed in specialized host cells called bacteriocytes and are reliably vertically transmitted from mother to offspring (2, 13). The obligate symbiont, *Buchnera aphidicola*, plays an essential role producing nutrients lacking in the aphid diet. The relationship between *Buchnera* and its host epitomizes a common dynamic in natural symbioses in which bacteria and host are completely dependent on one another for survival. Aphids can also be associated with facultative symbionts such as *Candidatus* “Hamiltonella defensa”, *Candidatus* “Regiella insecticola”, or *Serratia symbiotica* (14). These facultative symbionts often provide advantages to their aphid hosts—such as increasing their thermotolerance or protecting them from parasitoid wasps—but are not essential for aphid survival (15, 16).

In 2011 a culturable strain of *Serratia symbiotica*, CWBI-2.3^T^, was isolated from the black bean aphid, *Aphis fabae* (17). CWBI-2.3^T^ is a member of a widespread clade of *S. symbiotica* that is distinct from the bacteriocyte-associated symbiont strains. These gut-associated *S. symbiotica* appear to be at a primitive or transitional stage of symbiosis with aphids (18, 19). More specialized facultative symbiont strains of *S. symbiotica*, such as Tucson and IS, are found primarily in bacteriocytes and the insect hemolymph and are faithfully transmitted to all of a colonized mother’s offspring (20). In contrast, *S. symbiotica* CWBI-2.3^T^ primarily colonizes the digestive tracts of aphids and exhibits only sporadic transmission to progeny. Support for CWBI-2.3^T^ being a transitional stage also comes from its genome, which is intermediate in size between genomes of the strictly symbiotic strains, and genomes of related free-living *Serratia* species (21, 22).CWBI-2.3^T^ is thought to spread in natural populations of aphids through two horizontal transmission routes. It is excreted in aphid honeydew (liquid feces) onto plant surfaces, which could lead to environmental transmission (18, 19, 23), and it has been reported to spread between aphids feeding on the same plant by colonizing the phloem (24).

We examined the potential of *Serratia symbiotica* CWBI-2.3^T^ for paratransgenesis in aphids. We found that many plasmids, gene expression parts, and techniques used in *Escherichia coli* function in CWBI-2.3^T^. We used a fluorescently labeled strain to show that *S. symbiotica* CWBI-2.3^T^ can be reliably established in the guts of multiple species of aphids by feeding. However, colonization with CWBI-2.3^T^ eventually leads to decreased survival of some species. Finally, we show that we can achieve inducible expression of GFP in engineered CWBI-2.3^T^ living inside the aphid gut. No other aphid symbionts have been genetically engineered to date. Thus, CWBI-2.3^T^ represents a promising chassis organism for achieving short-term paratransgenesis, as well as an intriguing organism in which the genetic approaches that we describe could be used to study evolutionary transitions from insect pathogens to symbionts.

## RESULTS

### Genetic engineering of *S. symbiotica* CWBI-2.3^T^

We first tested whether common genetic techniques and DNA parts functioned in *S. symbiotica* CWBI-2.3^T^. We began by measuring the growth rate of the wild-type strain. Its doubling time in culture is approximately 4 hours. Accounting for this slower growth rate allowed for the successful transformation of CWBI-2.3^T^ through conjugation and electroporation procedures used in *E. coli*. By conjugating a nonreplicating donor plasmid encoding a mini-Tn7 construct from *E. coli* into *S. symbiotica* CWBI-2.3^T^, we were also able to integrate GFP into a specific location in its chromosome. Integration ensures more stable expression of engineered constructs and enables their functions to be maintained over many days without the need for antibiotic selection. This GFP-expressing *S. symbiotica* CWBI-2.3^T^ strain (CWBI-2.3^T^-GFP) can be used to monitor colonization of insects.

We also screened CWBI-2.3^T^ for its ability to maintain plasmids containing different origins of replication and antibiotic resistance cassettes, some of which have been reported to function in *Serratia marcescens* strains (25–28). We transformed plasmids that had either broad-host-range or *E. coli*-specific origins, as well as different copy numbers. Additionally, we tested whether CWBI-2.3^T^ was compatible with several common antibiotic resistance genes. We achieved successful transformants for every plasmid and antibiotic tested (Table 1). CWBI-2.3^T^ transformed with pBTK570, an RSF1010 plasmid with spectinomycin resistance that expresses E2 Crimson is shown in Figure 1A. These results demonstrate great flexibility in how one can genetically modify CWBI-2.3^T^ using combinations of plasmids and genome integration.

**FIG 1.**
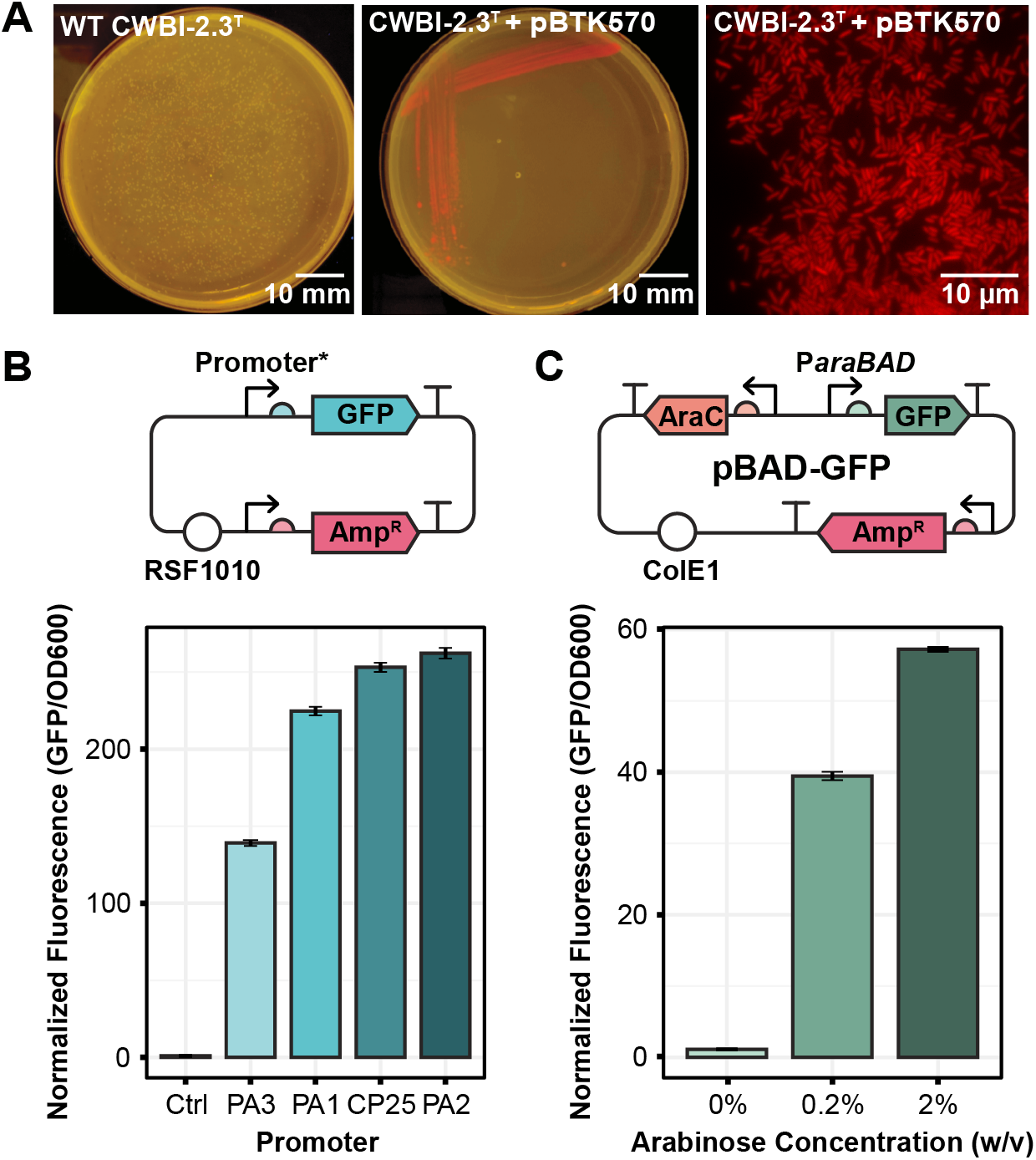
Origin and promoter function in *Serratia symbiotica* CWBI-2.3^T^. (A) Representative images of CWBI-2.3^T^ transformed with an RSF1010 origin plasmid expressing E2 Crimson. The left two images comparing this strain to wild-type CWBI-2.3^T^ were captured at the macro scale using a blue light transinducer and edited with linear adjustments. The image on the right was captured at the micro scale using a Nikon Eclipse inverted fluorescence microscope (640 nm excitation, 685 nm emission) and was linearly adjusted and pseudocolored. (B) Normalized expression of GFP from a series of plasmids that are identical except for their GFP promoter sequences. The plasmid map is shown above and expression levels are shown below. Expression for each promoter is normalized to the GFP/OD600 reading for wild-type CWBI-2.3^T^ (Ctrl). (C) Normalized expression of GFP from CWBI-2.3^T^-pBAD-GFP following induction with arabinose. A schematic of pBAD-GFP is shown above and expression levels are shown below. Fluorescence at each inducer concentration is normalized to the GFP/OD600 reading for wild-type CWBI-2.3^T^.

**Table 1.** Plasmid origins and antibiotic resistance genes that function in *S. symbiotica* CWBI-2.3T

| Origin | Copy number | Host range | Plasmid(s) | Reference |
| --- | --- | --- | --- | --- |
| RSF1010 | Low | Broad | pBTK570, pBTK501, pBTK503, pBTK509, pBTK510 | (28) |
| p15a | Low | Narrow | pBTK409 | (28) |
| pBBR1 | Medium | Broad | pAK*gfplux1* | (29) |
| ColE1 | High | Narrow | pYTK001, mTagBFP2-pBAD | (30, 31) |
| Antibiotic resistance | Resistance gene | Construct(s) |  | Reference |
| Ampicillin/Carbenicillin | *bla*<sub>TEM-116</sub> | pAK*gfplux1*, pBTK409, pBTK501, pBTK503, pBTK509, pBTK510 |  | (28, 29) |
| Ampicillin/Carbenicillin | *bla*<sub>TEM-181</sub> | mTagBFP2-pBAD |  | (31) |
| Chloramphenicol | *catI* | pYTK001 |  | (30) |
| Gentamicin | *aacC1* | pBT20 |  | (32, 33) |
| Kanamycin/Neomycin | *aphA-2 (nptII)* | miniTn7-GFP-Kan |  | (28, 34) |
| Spectinomycin | *aadA* | pBTK570 |  | (28) |

To complete our assessment of the capabilities of *S. symbiotica* CWBI-2.3^T^ we tested the performance of different constitutive and inducible promoters. For applications requiring continuous gene expression, we examined the relative levels of GFP expression from four broad-host-range constitutive promoters: PA1, PA2, PA3, and CP25 (28). Each of these promoters was functional, yielding a fluorescent signal 100-200 times brighter than the background autofluorescence of the wild-type strain (Fig. 1B). To allow for controlled gene expression, we also tested the functionality of the pBAD plasmid system, which uses the *araC* regulator to enable arabinose-inducible expression from the *araBAD* promoter. We found that this system behaves as expected. There was low expression of GFP in the absence of arabinose: the average fluorescence of the uninduced strain containing the pBAD-GFP plasmid was only 1.12 times the background of wild-type CWBI-2.3^T^ with no plasmid. GFP fluorescence in the strain with the pBAD-GFP plasmid became 39 and 57 times brighter than the wild-type strain when induced with 0.2% or 2% arabinose, respectively (Fig. 1C).

### Engineered *S. symbiotica* CWBI-2.3^T^ can colonize multiple species of aphids

To test the versatility of engineered CWBI-2.3^T^ for paratransgenesis we assessed its ability to colonize multiple species of aphids and monitored the colonization dynamics using CWBI-2.3^T^-GFP. We first established a method for delivering engineered CWBI-2.3^T^ to aphids by adapting the methods of Renoz *et al*. (23). CWBI-2.3^T^-GFP was delivered to 3^rd^ instar aphids through feeding on an artificial diet. Three days after feeding, a blue light transilluminator could be used to observe GFP fluorescence from CWBI-2.3^T^-GFP inside the colonized aphids. Using this feeding technique with *Ac. pisum*, we were able to reliably colonize an average of about 75% of treated aphids.

The fluorescence of the CWBI-2.3^T^-GFP strain also allowed us to monitor how CWBI-2.3^T^ colonizes living aphids. We observed that over the course of 10 days *S. symbiotica* CWBI-2.3^T^ spreads through the digestive tract of *Ac. pisum* until the entire gut is colonized (Fig. 2A). No bacteria were observed outside of the gut, and the gut remained colonized for the remainder of each aphid’s life. We also quantified host colonization by tracking the number of colony-forming units (CFUs) per aphid. On 2,3,5 and 10 days post-feeding, colonized *Ac. pisum* were crushed and plated to determine the bacterial load in their bodies. CWBI-2.3^T^ titer increased over time until it reached an average of about 10^8^ CFUs per aphid on day 5. Between day 5 and day 10, no significant change in titer was observed (*p* = 0.7529, two-tailed *t*-test) (Fig. 2B). All of these results are consistent with prior studies that used fluorescence *in situ* hybridization and quantitative PCR to monitor *Ac. pisum* colonization (23).

**FIG 2.**
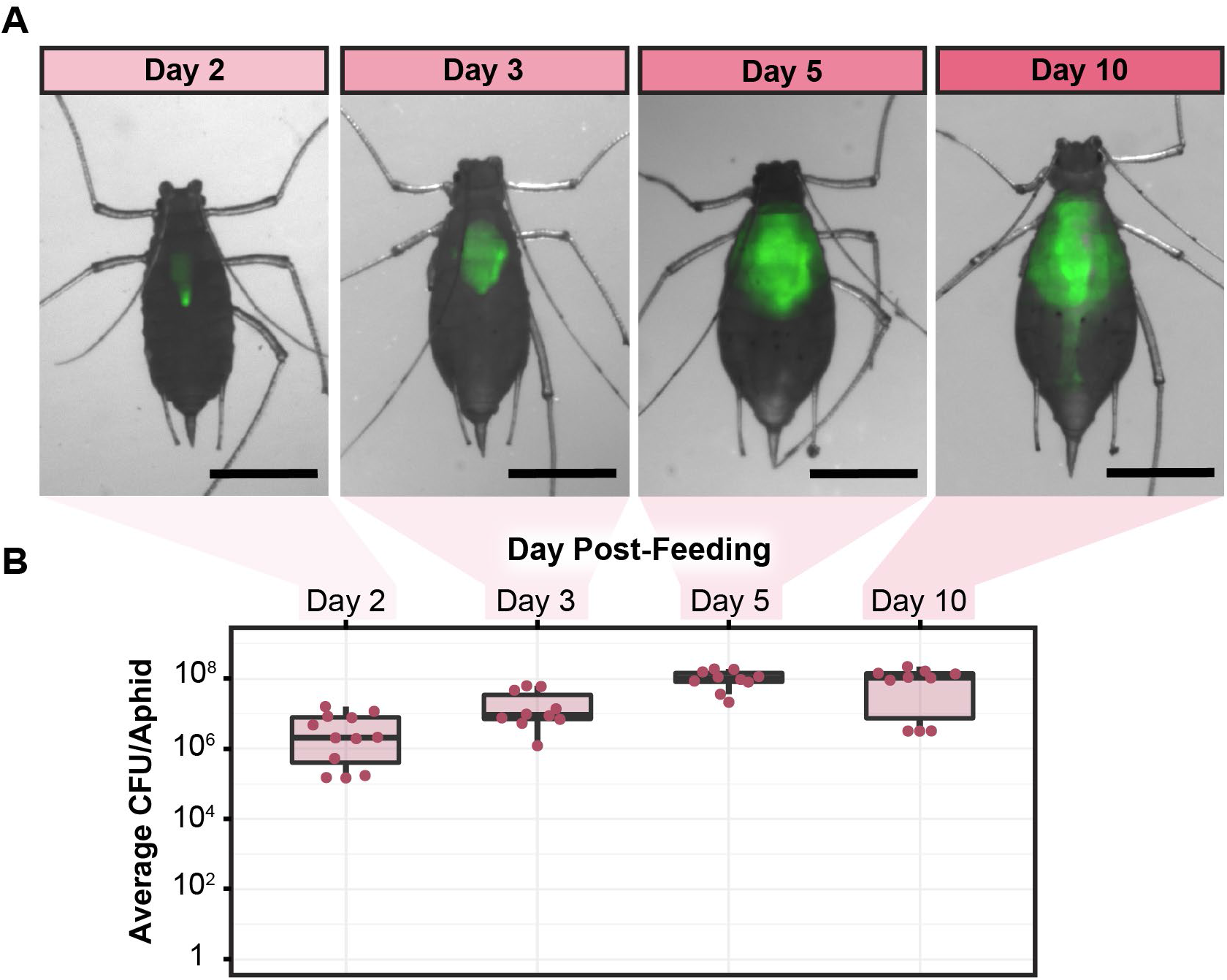
Colonization dynamics of CWBI-2.3^T^ in *Ac. pisum*. (A) Composite brightfield and fluorescence images tracking the spread of CWBI-2.3^T^-GFP through the gut of a live *Ac. pisum* individual. Dorsal images of the same aphid were captured for each time point. Images were linearly adjusted to enhance the brightness of GFP. Scale bar is 1 mm. (B) CWBI-2.3^T^-GFP titer in *Ac. pisum* over time. At least 10 aphids per time point were crushed and plated to perform CFU counts.

We next tested whether the colonization pattern in *Ac. pisum* was representative of other aphid species by repeating this experiment with *Aphis fabae* (black bean aphid), *Aphis craccivora* (cowpea aphid), and *Lipaphis erysimi* (mustard aphid). We found that CWBI-2.3^T^-GFP principally colonizes the gut of these three species as well (Fig. 3A). Each of the other aphids tested also showed a similar trend in bacterial titer over time. The CFUs per aphid consistently reached between 10^7^ and 10^8^ on the fifth day after treatment for all four species, with averages of 5.3 × 10^7^, 7.4 × 10^7^, 2.4 × 10^7^, and 1.2 × 10^7^ for *Ac. pisum, A. fabae, A. craccivora*, and *L. erysimi*, respectively (Fig. 3B). There were slight, but significant differences in bacterial titers observed in the four species (*F*_3,32_ = 6.792, *p* = 0.00113, one-way ANOVA). Titer was highest in *A. fabae*, which is the natural host from which CWBI-2.3^T^ was isolated. The trend in the other species may reflect their different body sizes, as *Ac. pisum* is much larger than the other aphids.

**FIG 3.**
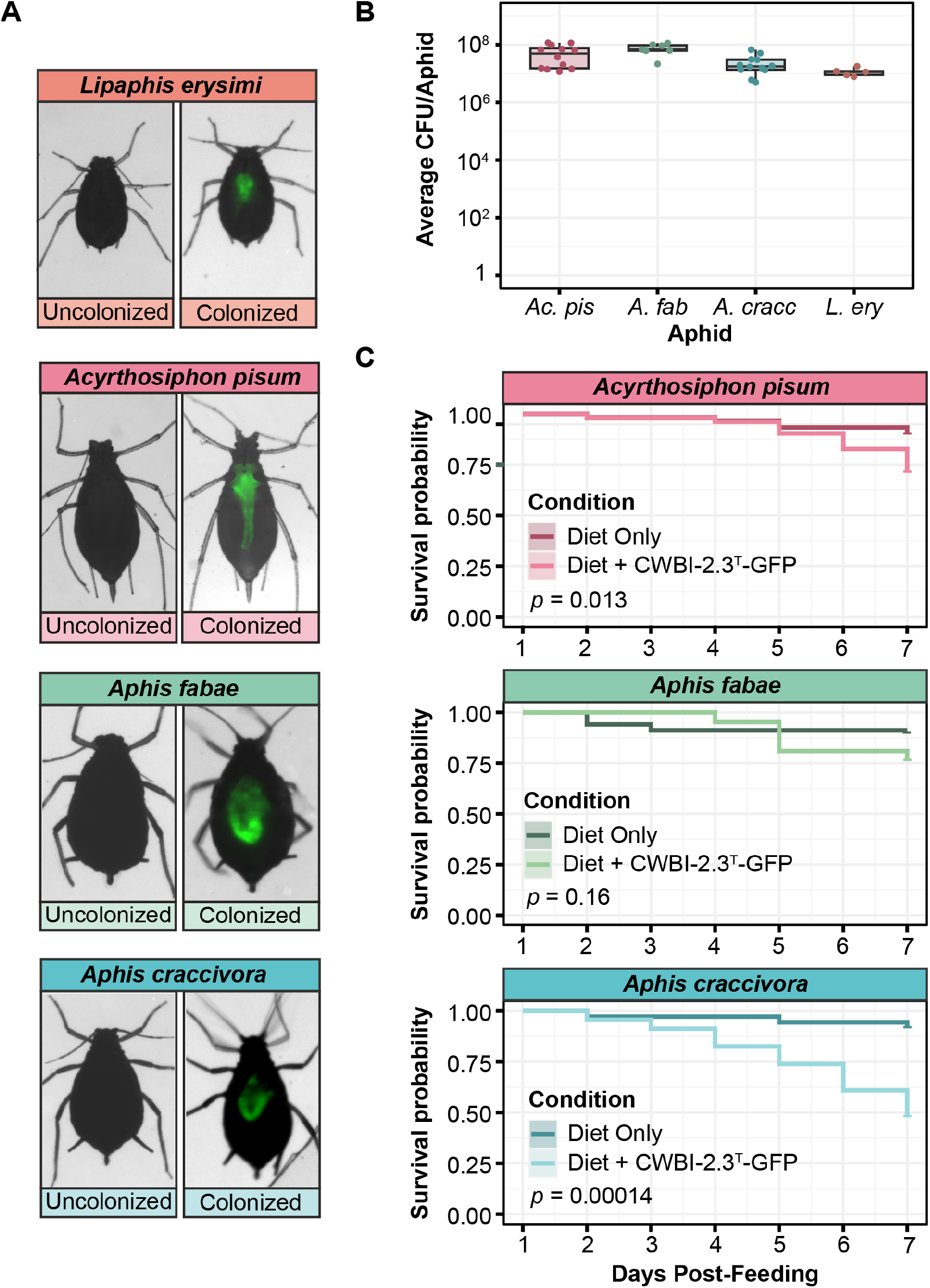
CWBI-2.3^T^-GFP colonization of multiple aphid species. (A) Composite brightfield and fluorescent stereoscope images showing CWBI-2.3^T^-GFP colonization of four different species. Photographs for the uncolonized and colonized pairs were captured with the same exposure for comparison. Images were captured from the ventral side of each aphid on the 5^th^ day after feeding. (B) CWBI-2.3^T^-GFP titer in each of the 4 species on day 5 post-feeding. At least 5 aphids per species were crushed and plated to perform CFU counts. (C) Mortality curves showing the survival probability of each aphid colonized with CWBI-2.3^T^-GFP compared to uncolonized aphids. Counts began on the first day after feeding to eliminate aphids that died during experimental setup. The starting population was at least 21 aphids for each survival curve. The statistical significance of differences in aphid survival were calculated using a log-rank test (*p* values shown in each panel).

### *S. symbiotica* CWBI-2.3^T^ decreases the survival of some aphid hosts

Next, we investigated how CWBI-2.3^T^ affected a component of aphid fitness. We monitored the survival of aphids for 7 days after colonizing them with CWBI-2.3^T^-GFP through feeding (Fig. 3C). We observed a similar impact of CWBI-2.3^T^-GFP on the survival of *Ac. pisum* and *A. fabae*. After 7 days, the treated *A. fabae* and *Ac. pisum* populations lost 15% and 19% more aphids than their respective control populations. This decrease was not significant for *A. fabae* and only marginally significant for *Ac. pisum (p* = 0.16 and *p* = 0.013, log-rank tests, respectively). These results are consistent with previous studies that used wild-type CWBI-2.3^T^ (19, 23). Colonization with engineered CWBI-2.3^T^ had a larger effect on *A. craccivora* survival. After 7 days, 44% fewer colonized *A. craccivora* remained alive compared to the uncolonized aphids of this species, and this reduction was highly significant (*p =* 0.00014, log-rank test). The effects of CWBI-2.3^T^ colonization on *A. craccivora* have not been tested before. Overall, these results show that CWBI-2.3^T^ can have a range of fitness effects on different aphid host species.

### Induction of *S. symbiotica* CWBI-2.3^T^ can be performed *in vivo*

To enable the controlled expression of genes inside living aphids, we tested whether we could induce the expression of GFP inside *A. pisum*. We first colonized *Ac. pisum* with wild-type CWBI-2.3^T^ or CWBI-2.3^T^ transformed with the plasmid pBAD-GFP (strain CWBI-2.3^T^-pBAD-GFP). Three days later we fed those same aphids on leaves embedded in agar containing no arabinose or 2% arabinose. We observed that after one day of feeding, only the aphids that were colonized with CWBI-2.3^T^-pBAD-GFP and fed on 2% arabinose showed visible GFP fluorescence (Fig. 4A). We quantified the per-aphid fluorescence in each condition and found that arabinose induction significantly increased the GFP signal in the CWBI-2.3^T^-pBAD-GFP aphids (*p* = 6.9 × 10^−8^, one-tailed Mann-Whitney U test) (Fig. 4B). On average, GFP fluorescence was 83.5 times brighter with induction compared to the uninduced condition.

**FIG 4.**
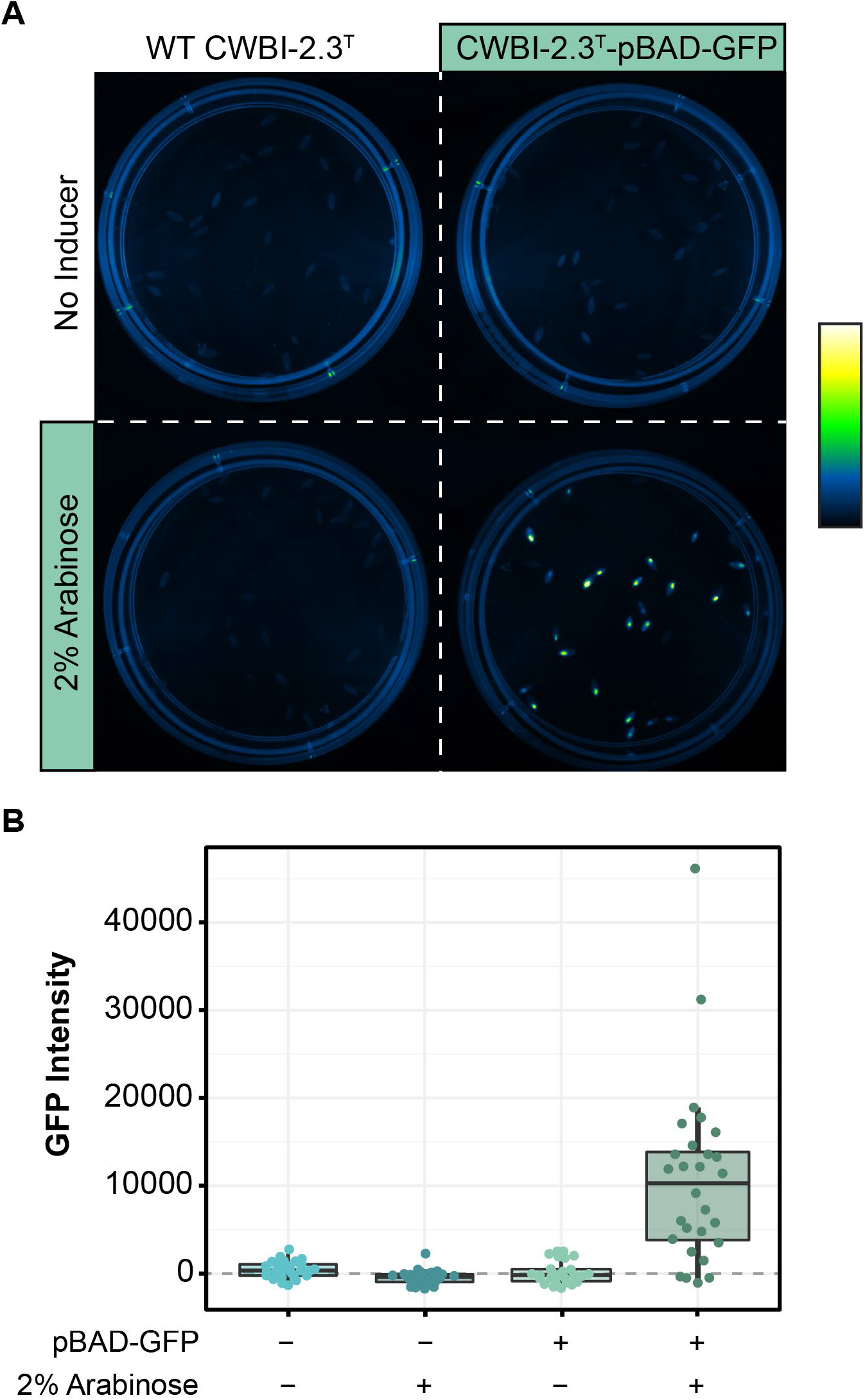
GFP induction inside living aphids. (A) Photograph of GFP expression for each of the four conditions tested for induction: CWBI-2.3^T^ ± Arabinose and CWBI-2.3^T^-pBAD-GFP ± Arabinose. The image has been pseudocolored and linearly adjusted for contrast. (B) Boxplots showing the fluorescence distribution of each population of treated aphids. The GFP intensity of at least 24 aphids was measured per condition.

We confirmed that the lack of fluorescence in the uninduced CWBI-2.3^T^-pBAD-GFP condition was not due to poor colonization or loss of plasmid function by crushing the aphids and plating their bacteria on media containing 2% arabinose. We found that 93% and 100% of the aphids in the uninduced and induced CWBI-2.3^T^-pBAD-GFP conditions, respectively, were colonized with *S. symbiotica*. In both conditions, the plated bacteria all expressed GFP, indicating that the plasmid had not been lost or mutated. The discrepancy between the percentage of aphids that were colonized (100%) and how many visibly fluoresced (82.8%) in the induced condition and variation in the fluorescence per aphid are likely attributable to differences in when and how much individual aphids fed on the inducer under the experimental conditions.

## DISCUSSION

In this study we establish a platform that can be used for aphid paratransgenesis. We found that *Serratia symbiotica* CWBI-2.3^T^ is compatible with both broad-host-range and *E. coli* plasmids and gene expression parts. We fed engineered CWBI-2.3^T^ back to multiple aphid species, tracked gut colonization over time, and characterized the effects on insect survival. Lastly, we showed that an engineered function in CWBI-2.3^T^ could be induced within living aphids. This work provides new tools that can be used to increase our understanding of aphid/symbiont relationships, as well as a foundation for developing methods for the control of aphids pests without the need for insect or plant genetic engineering.

Previous studies of CWBI-2.3^T^ characterized its relationship with *A. fabae* and *Ac. pisum* (19, 23). These experiments established the basis for the hypothesis that CWBI-2.3^T^ and related strains may resemble a transitional proto-symbiont of aphids. In particular, *S. symbiotica* CWBI-2.3^T^ does not colonize aphid bacteriocytes or exhibit vertical transmission to offspring (18, 19), which are important characteristics of obligate and facultative bacterial symbionts that have co-evolved with aphids. Our results with engineered CWBI-2.3^T^ support the original findings in *Ac. pisum* and *A. fabae*, and we find similar results in two additional aphid species: *A. craccivora* and *L. erysimi*. CWBI-2.3^T^ colonizes the guts of all these aphids and can have mild to moderate negative effects on aphid fitness. These results support the hypothesis that this strain represents a transitional state of symbiosis.

*S. symbiotica* provides a unique opportunity to study the evolution of aphid symbioses because the entire spectrum of known host-microbe relationships are represented within a single bacterial species (21, 22). CWBI-2.3^T^ and its relatives are aphid proto-symbionts and retain their ability to survive outside the host. *S. symbiotica* Tucson, IS, and similar strains are more specialized facultative symbionts of many aphid species. *S. symbiotica* SCt, SCc, and STs have evolved to act as co-obligate symbionts with *B. aphidicola* in Lachninae aphids. As “transitional” symbionts, CWBI-2.3^T^ and related *S. symbiotica* strains likely have genetic features that distinguish them from the more specialized endosymbiont clades. For example, the genome of the CWBI-2.3^T^, while reduced, is about 1 Mb larger than that of the bacteriocyte-associated *S. symbiotica* Tucson strain found in *Ac. pisum* (21). The genomes of the CWBI-2.3^T^-like *S. symbiotica* strains encode potential pathogenicity factors that are missing in the bacteriocyte-associated strains, which could contribute to their more volatile relationships with their hosts (35).

We expect that our work will empower future studies of the proto-symbiont relationship between CWBI-2.3^T^ and aphids. By engineering CWBI-2.3^T^ we were able to observe colonization dynamics in living aphids and easily distinguish the colonization status of the aphids due to their fluorescence. This feature simplifies large assays where colonization is required, such as mortality or transmission experiments. Similar work engineering *Arsenophonus nasoniae*, a parasitoid wasp symbiont, to express GFP allowed researchers to better understand how it is vertically transmitted during oviposition (36). In the future, the ability to engineer CWBI-2.3^T^ and deliver it back to aphids could be combined with engineering tools for gene knockout. This approach was used to study the role of the type III secretion system of *Sodalis glossinidius* in establishing a symbiotic relationship with its tsetse fly host (37). Similar genetic approaches could be used to understand the role of putative symbiotic factors in CWBI-2.3^T^ that may have enabled its transition from a free-living, plant-associated bacterium to an insect symbiont.

Engineering CWBI-2.3^T^ to manipulate aphid biology could also be of interest for agricultural applications. It has been proposed that insect paratransgenesis could be used for targeted pest-control strategies that pose less of a risk to the environment than chemical pesticides (9, 38). In aphids, one approach would be to engineer CWBI-2.3^T^ to reduce the capacity of aphids for vectoring plant diseases. Related techniques have already been carried out by groups performing paratransgenesis in other insects. For instance, *Pantoea agglomerans*, a symbiont of the glassy-winged sharpshooter, was engineered to produce antimicrobial peptides that selectively kill the phytopathogen *Xylella fastidiosa* that is carried and spread by sharpshooters (8). Another *Serratia* symbiont, *Serratia* AS1, was engineered to limit mosquito transmission of the malaria parasite (5). These approaches took advantage of the fact that both symbiont and pathogen occupied the same gut environment within their host to successfully reduce pathogen transmission. In aphids, CWBI-2.3^T^ could be used to similarly prevent the spread of bacterial phytopathogens that propagate within aphids such as *Erwinia aphidicola*, *Dickeya dadantii*, and *Pseudomonas syringae* (39).

Our ability to engineer CWBI-2.3^T^ could also be adapted to enable symbiont-mediated RNA interference (RNAi) to control aphid gene expression and vectorial capacity. This method enables targeted gene silencing by engineering the symbiont to produce double-stranded RNA inside the host to induce its innate RNAi response. Symbiont-mediated RNAi has proven to be effective for genetically manipulating insects, such as kissing bugs and western flower thrips, that are typically less compatible with injecting or feeding double-stranded RNA (4, 6). This approach has also recently been used to protect honeybees from parasitic mites and deformed wing virus (40). In aphids, symbiont-mediated RNAi could be used to directly target plant viruses that circulate and/or propagate in aphids (e.g., Luteoviruses such as potato leaf roll virus and Rhabdoviruses such as lettuce necrotic yellows virus) (41, 42). It might also be used to reduce the expression of the receptors that noncirculative viruses bind to such as the Stylin-01 receptor bound by cauliflower mosaic virus (43).

Overall, the capabilities for engineering *S. symbiotica* CWBI-2.3^T^ that we have demonstrated could lead to a multitude of new applications in the future. They may help researchers build a better understanding of the evolution of host-microbe symbioses in the well-established aphid model system. These tools also provide a foundation for exploring new synthetic biology approaches for pest management that could lead to safer and more environmentally friendly agricultural practices.

## MATERIALS AND METHODS

### Growth of *S. symbiotica* CWBI 2.3^T^

We obtained *S. symbiotica* CWBI 2.3^T^ from the German Collection of Microorganisms and Cell Cultures (DSM 23270). Bacteria were cultured at room temperature (~25°C) in BBL Trypticase Soy Broth (TSB) or on Trypticase Soy Agar (TSA) plates (Becton, Dickinson and Company, MD, USA). For liquid cultures, the tubes were incubated with orbital shaking at 180 r.p.m. over a diameter of one inch. Concentrations of antibiotics and other media supplements used in this study are as follows: carbenicillin (100 μg/mL), chloramphenicol (20 μg/mL), gentamicin (40 μg/mL), kanamycin (50 μg/mL), spectinomycin (60 μg/mL), and diaminopimelic acid (DAP)(0.3 mM). To assess growth rate, two-day-old cultures of CWBI-2.3^T^ were grown up and diluted 1:100 into fresh TSA in a 96 well plate. Ten replicates were included on the plate which was incubated at 25°C with 6 mm amplitude orbital shaking every 15 seconds in an Infinite 200 Pro plate reader (Tecan, Männedorf, Switzerland). OD600 readings were taken every 10 minutes for 48 hours. Growth curves were fit to a logistic model using Growthcurver (version 0.3.0) (44).

### Growth and maintenance of aphid colonies

*Acyrthosiphon pisum* LSR1 was acquired from long-term stocks maintained in the lab of Nancy Moran (University of Texas at Austin). *Aphis fabae* was obtained from the lab of Thierrey Hance (Université catholique de Louvain, Belgium). *Aphis craccivora* and *Lypaphis erysimi* were collected in Austin, TX and identified by COI barcode sequencing using the LepF (5’-ATTCAACCAATCATAAAGATATTGG-3’) and LepR (5’-TAAACTTCTGGATGTCCAAAAAATCA-3’) primer pair (45). *Ac. pisum, A. fabae*, and *A. craccivora* were maintained on Broad Windsor *Vicia faba* plants (Mountain Valley Seed Company, UT, USA). *L. erysimi* was reared on *Brassica oleracea* var. Capitata (The Seed Plant, TX, USA). All colonies were maintained in cup cages on their respective plants at 20°C with a long (16L:8D) photoperiod in Percival I-36LLVL incubators (Perry, IA, USA).

### Transformation of *S. symbiotica* CWBI 2.3^T^

*S. symbiotica* CWBI 2.3^T^ was made electrocompetent using a modification of a protocol for *E. coli*. Cultures were first grown for two days until they reached saturation. Fifty microliters of saturated culture were then inoculated into 50 mL of TSB in a 250 mL flask and grown for about 16 h to mid-log phase (OD600 = 0.4-0.6). Cells were then centrifuged at 4500 × *g* for 5 min, the supernatant was discarded, and the cells were resuspended in 40 mL of 10% glycerol. This wash step was repeated 4 additional times. The pellet was then resuspended in 500 μL of 10% glycerol, divided into 50 μL aliquots, and frozen at −80°C. For electroporation, 2 μL plasmid was added to 50 μL of electrocompetent cells and electroporated at 2.5 V in a 0.1 cm cuvette with the BioRad MicroPulser (Hercules, CA, USA). The cells were resuspended in 950 μL TSB and allowed to recover overnight (~16 h) then plated on TSB agar with selective antibiotic.

For conjugative transformation, cultures of donor *E. coli* strain, MFDpir and the recipient *S. symbiotica* CWBI-2.3^T^ strain were first grown to saturation. Then, 1 mL of each was centrifuged at 1000 × *g* for 5 min. The supernatant was discarded, and the pellets were resuspended in 1 mL 145 mM saline and centrifuged again as before. The wash step was repeated again, and both pellets were resuspended in 500 μL of 145 mM saline. A 50 μL 1:100 ratio of donor:recipient cells was prepared in a separate Eppendorf tube then spot plated on a TSB + DAP plate. After two days of growth on the conjugation plate, a metal loop was used to scrape up a small pellet of bacteria from each spot and place it into 1 mL of saline solution. The pellets were then centrifuged and washed twice with saline as described in the steps for the previous day. Next, 100 μL of resuspended solution along with 100 μL of 10× concentrated solution was plated on TSA plates containing selective antibiotics. After two to three days of growth, successfully conjugated CWBI-2.3^T^ colonies were picked and screened for proper strain identity and presence of the conjugative plasmid.

### Mini-Tn7 integration into the CWBI-2.3^T^ chromosome

Integration of a GFP and kanamycin resistance gene cassette into *S. symbiotica* to create strain CWBI-2.3^T^-GFP was carried out using the mini-Tn7 system (34). Integration was performed according to the conjugation steps described above, with modifications for the use of two MFDpir^+^ donor strains. One donor strain expresses the suicide delivery vector containing the genes of interest (GFP/Kan^R^), and the other holds the helper plasmid (pTNS2) containing the components of the TnsABCD site-specific transposition pathway. Conjugation of both plasmids should insert the GFP/Kan^R^ genes into the chromosome of *S. symbiotica* 25 base pairs downstream of the *glmS* gene at the *att*Tn7 site. Chromosomal integration at the expected site was confirmed through PCR and Sanger sequencing using the GFP-mut3-forward (5’-AGCCGTGACAAACTCAAGAA-3’) and *glmS*-reverse (5’-GCCGTTGCAATTGTTGTC-3’) primer pair.

### Assessing plasmid origin, antibiotic resistance gene, and promoter function in CWBI-2.3^T^

The compatibility of different origins of replication and antibiotic resistance genes was tested by transforming CWBI-2.3^T^ with various plasmids, as shown in Table 1. Transformants were picked, and plasmids were purified from cells and verified by Sanger sequencing to confirm successful transformation.

Gene expression from various synthetic promoters was tested by transforming pBTK501, pBTK503, pBTK509, and pBTK510 from the bee microbiome toolkit into CWBI-2.3^T^ (28). Each strain was grown in culture for two days, then transferred to a fresh tube and diluted to an OD of 0.05 in triplicate. Following an additional day of growth, GFP fluorescence was measured in the Tecan Infinite 200 Pro plate reader (485 nm excitation, 535 nm emission). Three technical replicates were measured for each of the three biological replicates.

To construct the pBAD-GFP plasmid, we used Gibson assembly to combine the pBR322 origin, *araC* regulator, *araBAD* promoter, and ampicillin resistance gene from the mTagBFP2-pBAD plasmid (Addgene #54572) with the GFPmut3 gene from pBTK503 (28, 31). Overnight cultures of CWBI-2.3^T^ transformed with pBAD-GFP (CWBI-2.3^T^-pBAD-GFP) were spiked with 0.2 or 2% (w/v) arabinose. Three biological replicates of each condition, including the no arabinose control, were used for this experiment. After 24 additional hours of growth, GFP fluorescence was measured using the Infinite 200 Pro plate reader (485 nm excitation, 525 nm emission).

### Feeding *S. symbiotica* CWBI-2.3^T^ to aphids

Feeding was carried out using age-controlled aphid populations reared on *V. faba*. Third instar aphids were used for all feeding experiments. Cultures of *S. symbiotica* CWBI-2.3^T^ were first grown to log phase, centrifuged at 1000 × *g* for 5 min, washed twice with 1× PBS, then resuspended in 1× PBS to an OD of 1. One microliter of this culture, along with 2 μL of yellow food dye to improve aphid feeding rates (Gel Spice Company Inc., NJ, USA), was added per 100 μL of Febvay artificial diet (46). Feeding chambers were assembled using 33 mm petri dishes and two layers of parafilm. A hole was cut into the bottom of one half of the petri dish and covered with a piece of tightly woven mesh for air flow. A single piece of parafilm was stretched thin over the other half, and 100 μL of diet was pipetted onto the parafilm. The other piece of parafilm was stretched thin over top of the diet to create a “sandwich”. Up to 30 aphids were added to the other half of the petri dish and both sides were sealed together with parafilm. Aphids were fed for 16-24 h then transferred to *V. faba* plants and observed as needed.

### Imaging aphid colonization

Five-to six-day-old aphids were fed on diet plates containing CWBI-2.3^T^-GFP as described in the quantification methods above. For the time-course, *Ac. pisum* was checked for colonization on the second day post-feeding using a GBox-F3 Syngene imager, and 4 were selected to be imaged for the time points at 2, 3, 5, and 10 days post-treatment. Between all imaging time points the aphids were maintained on separate *V. faba* plants under normal conditions. For the images of the different aphid species at one time point, aphids were screened for colonization on day 5 and 3 of each were randomly selected to be imaged. A Leica MZ16 Fluorescent Stereoscope was used in conjunction with the Leica Application Suite software to capture all images. Images were captured in virtual stacks using both a brightfield and GFP channel. To qualitatively highlight the localization of the colonizing bacteria, the brightness and contrast of the images were linearly adjusted using ImageJ (version 1.52p) as necessary.

### Quantifying colonization of aphids with CWBI-2.3^T^

Each species of aphid was first fed on diet containing CWBI-2.3^T^-GFP at 5-6 days of age as described above. Aphids were fed in pools of 25-30 aphids per container. After 24 hours, aphids were moved to *V. faba* plants. The day when the aphids were transferred to plants serves as day 0 for the colonization experiments. On days 2, 3, 5, and 10, aphids were collected and processed. First, aphids were surface sterilized by soaking in 200 μL 10% bleach for 1 minute. Aphids were then rinsed with distilled water, resuspended in 200 μL saline and squashed with a pestle. The squashed aphid mixture was serially diluted 10-fold to reach final 10^5^ dilution. Five microliters of each dilution mix were spotted onto TSA + kanamycin plates (3 replicates per aphid). The plates were incubated for 3 days and then colonies were counted to determine the CFU per aphid.

### Aphid fitness following colonization with CWBI-2.3^T^

We assessed the effect of CWBI-2.3^T^ on the fitness of each aphid species by measuring mortality rates. Aphids were fed on either plain diet or diet containing CWBI-2.3^T^-GFP. Following treatment, they were transferred to plants, and the number of living adult aphids was recorded each following day. To determine whether aphids that died prior to day 5 were colonized with CWBI-2.3^T^ (before it expressed sufficient GFP to be detected by eye), these dead aphids were collected, washed in bleach, squashed and plated as described in the quantification section above. Pieces of mesh material were wrapped around the base of each plant at the start of the experiment so that dead aphids could be easily collected. On day 5, the remainder of the uncolonized aphids were visually sorted and removed from the experimental pool. Mortality was recorded for a total of 7 days post-treatment.

### Inducible control of GFP expression *in vivo*

The assay for the function of *in vivo* induction was carried out in two separate feeding steps: colonization with CWBI-2.3^T^ and induction with arabinose. To colonize the aphids with the symbiont, aphids were first fed on diet plates containing either wild-type CWBI-2.3^T^ or CWBI-2.3^T^-pBAD-GFP. After feeding on diet for 24 hours the aphids were transferred to fava beans. Induction plates were created using 2 cm diameter petri dishes. Each dish was propped up at an angle, and 1 mL 1.5% agar with or without 2% (w/v) arabinose was added. *V. faba* leaves were added to the agar as it cooled, and plates were stored at 4°C. All induction plates were prepared 2 days before aphids fed on them. The aphids were transferred to the induction plates after resting on the fava bean plants for 3 days. Three plates containing 30 aphids each were set up for each condition: CWBI-2.3^T^, CWBI-2.3^T^ + 2% arabinose, CWBI-2.3^T^-pBAD-GFP, and CWBI-2.3^T^-pBAD-GFP + 2% arabinose.

After 24 hours of feeding on the induction plates, 30 aphids per condition were randomly selected and pooled, and GFP expression of all the aphids was photographed using the GBox-F3 Syngene imager using a blue LED excitation light and SW06 emission filter. Plates were photographed together in one image to ensure comparable fluorescence measures. To confirm that the aphids were colonized and that the number of aphids showing fluorescence was equivalent to the number of aphids colonized with CWBI-2.3^T^-pBAD-GFP, aphids from the induced and uninduced conditions were squashed and 10-fold dilutions were plated onto TSA + Carb + 2% (w/v) arabinose using the protocol used to quantify colonization of aphids described above.

Images were analyzed using ImageJ (version 1.52p). To measure GFP intensity the background was first subtracted from the image using the rolling ball method. Then, image intensity was linearly adjusted to bring the background value for the plate down to zero. This background subtraction step was performed separately for each plate in the image (i.e., each set of aphids in one condition). Finally, the GFP intensity for each aphid was determined by outlining it and using the measurement tool to calculate the integrated density within this region of interest. To account for aphid autoflourescence, the average integrated density of all aphids in both the wild-type CWBI-2.3^T^ conditions was subtracted from the measurements made in all four conditions.

## ACKNOWLEDGEMENTS

This work was supported by the Defense Advanced Research Projects Agency (HR0011-17-2-0052), the U.S. Army Research Office (W911NF-20-1-0195), and the University of Texas at Austin Stengl-Wyer Endowment. We thank François Renoz for assistance with acquiring aphids. We also thank Sean Leonard and Peng Geng for sharing plasmids and experimental advice. This research made use of equipment at the Microscopy and Imaging Facility of the Center for Biomedical Research Support at The University of Texas at Austin.

